# Coordinated electrical activity in the olfactory bulb gates the oscillatory entrainment of entorhinal networks in neonatal mice

**DOI:** 10.1101/352054

**Authors:** Sabine Gretenkord, Johanna K. Kostka, Henrike Hartung, Katja Watznauer, David Fleck, Angélica Minier-Toribio, Marc Spehr, Ileana L. Hanganu-Opatz

**Affiliations:** Developmental Neurophysiology, Institute of Neuroanatomy, University Medical Center Hamburg-Eppendorf, 20251 Hamburg, Germany; Department of Chemosensation, Institute of Biology II, RWTH Aachen University, 52074 Aachen, Germany

**Keywords:** Development, olfactory, entorhinal, oscillations, optogenetics, mitral cells, connectivity

## Abstract

While the developmental principles of sensory and cognitive processing have been extensively investigated, their synergy has been largely neglected. During early life, most sensory systems are still largely immature. As a notable exception, the olfactory system reaches full maturity during intrauterine life, controlling mother-offspring interactions and neonatal survival. Here, we elucidate the structural and functional principles underlying the communication between olfactory bulb (OB) and lateral entorhinal cortex (LEC) – the gatekeeper of limbic circuitry – during neonatal mouse development. Combining optogenetics, pharmacology, and electrophysiology *in vivo* with axonal tracing, we show that mitral cell-dependent discontinuous theta bursts in OB drive network oscillations and time the firing in LEC via axonal projections confined to upper cortical layers. Pharmacological silencing of OB activity diminishes entorhinal oscillations. Moreover, odor exposure boosts OB-entorhinal coupling at fast frequencies. Thus, early OB activity shapes the maturation of entorhinal circuits.

## INTRODUCTION

Coordinated patterns of electrical activity periodically entrain developing neuronal networks in rhythms with a broad frequency spectrum. These patterns have been proposed to critically shape brain maturation (1–3). Experimental evidence supporting this hypothesis has been mainly provided for sensory systems. For example, in the visual and auditory systems, spontaneous activity from sensory periphery (i.e., retina or cochlea) controls the formation of cortical representations underlying stimulus perception (4, 5). Theta band spindle bursts and gamma oscillations in the developing somatosensory system promote thalamo-cortical connectivity and maturation of coupling with the motor system (6, 7). Overall, the discontinuous oscillatory activity in sensory cortices during development has multifold origin, including stimulus-independent activation in the periphery and entrainment of local cortical circuits via chemical and electrical synapses (1, 8).

While less investigated, limbic circuits show similar patterns of coordinated activity during early development, with discontinuous theta bursts (4-12 Hz) and superimposed fast episodes (20-40 Hz) in beta-gamma frequency (9–13). Theta bursts facilitate unidirectional communication from the CA1 area of intermediate/ventral hippocampus (HP) to the prelimbic subdivision of the prefrontal cortex (PFC) via glutamatergic projections (14). As a consequence of hippocampal theta drive, pyramidal neurons in local prelimbic circuits generate beta-low gamma oscillations (15). Theta coupling between neonatal PFC and HP is controlled by the lateral entorhinal cortex (LEC) that densely projects to both areas (11). The complex organization of limbic circuits at early age raises the question, which mechanisms control the gatekeeper function of LEC during early development. Similar to sensory systems, the neonatal LEC could be driven by spontaneous activity from the sensory periphery. Indeed, the adult LEC receives direct input from the olfactory bulb (OB) that, in contrast to other sensory systems, bypasses the thalamus (16, 17). Mitral and tufted cells (MTCs) represent the sole OB output neurons. Rather than simply relaying information, these neurons are embedded in a complex network that controls odor information coding (18, 19). The axons of MTCs terminate in entorhinal layer I on apical dendrites of layer II/III pyramidal and stellate cells (20), which in turn form the perforant path projection to the hippocampal formation (21, 22). Layer II/III neurons in LEC project back to OB (23), yet distinct entorhinal populations are differently engaged in feedforward and feedback signaling during odor processing (24). Thereby, odor-evoked activity in the adult controls the gateway function of LEC interfacing HP and neocortical regions (25, 26).

While the sense of smell serves fundamental functions in newborn animals (27), the role of olfactory inputs and OB activity for limbic circuit maturation remains unknown. Since other sensory systems are still immature during early life – and thus their impact on limbic circuits is negligible – this knowledge gap appears even more striking. Rodent pups are blind, deaf and have limited sensorimotor abilities until the end of the second postnatal week (28, 29). In contrast, the olfactory system maturates early and is considered to be fully functional at birth, providing the major sensory stimulus in neonatal rodents. We hypothesize that both odor-dependent and -independent coordinated activity in OB control the entrainment of entorhinal networks during neonatal development. Here, we combine optogenetics, electrophysiology, and pharmacology *in vivo* with anatomical tracing in neonatal mice (postnatal day (P) 8-10) to elucidate the olfactory control of the functional maturation of entorhinal circuits.

## RESULTS

### OB and LEC are reciprocally connected in neonatal mice

In mice, MTCs mature during intrauterine life and their axons reach cortical targets during the first two postnatal weeks (30). This time window coincides with the period of strong gating of prefrontal-hippocampal networks by theta activity in LEC. To detail on the spatial patterns of connectivity between OB and LEC in P8-10 mice, we performed an in-depth investigation of axonal projections from MTCs to LEC and, *vice versa,* of entorhinal projections to OB. First, we used Tbet-cre;R26-tdTomato mice (n=4) for intact-brain imaging of long-range projections by electrophoretic tissue clearing and confocal fluorescence microscopy (Fig 1A, B). In these mice, MTCs are genetically tagged (Fig 1C) (31). Already at P8, the lateral olfactory tract (LOT) comprising MTC axons appeared fully developed and reached the posterior part of the cerebrum, including piriform cortex (PIR) and LEC (Fig 1A, D). As previously shown in adult rats (32), MTC axons were mainly confined to layer I of neonatal LEC (Fig 1D). Retrograde tracing with Fluorogold (FG) injected into the LEC of P3-4 mice confirmed the direct connectivity (Fig 1E). No differences between dorsal and ventral OB were detected with respect to the density of MTC projections to LEC.

**Fig 1.**
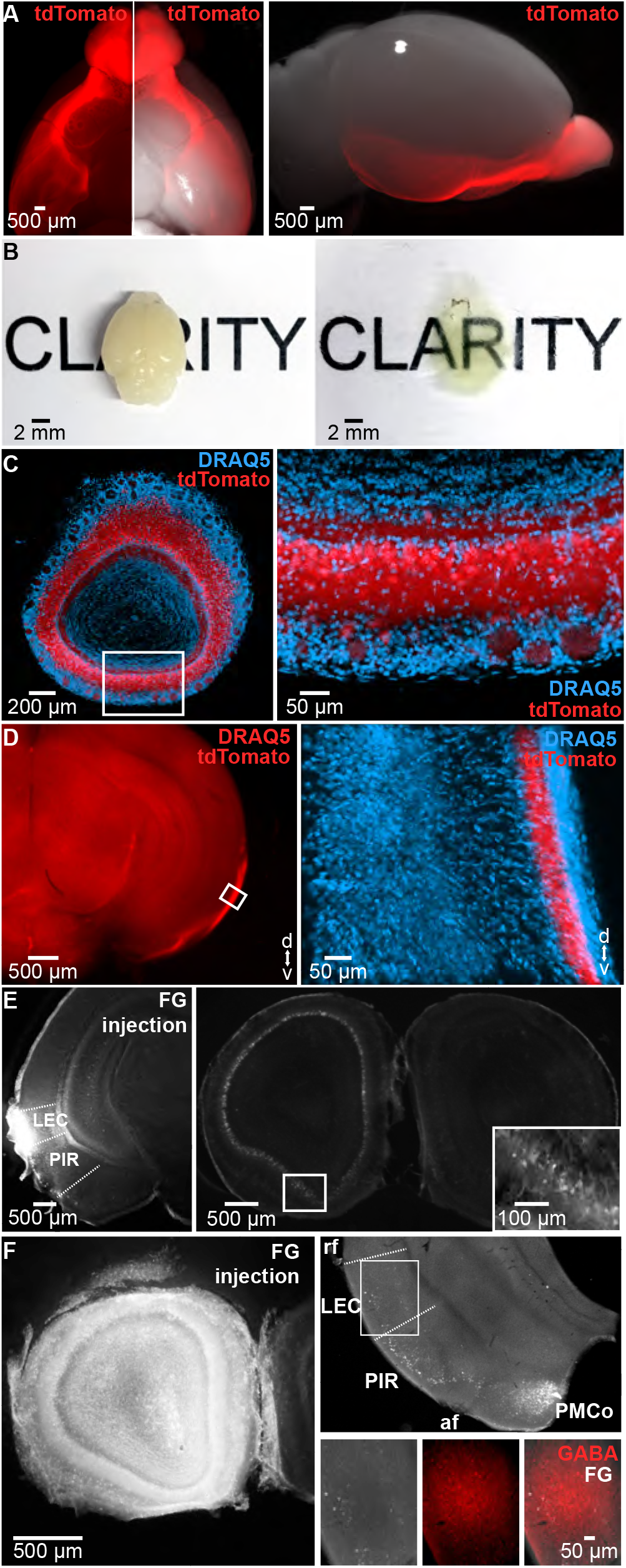
Patterns of axonal connectivity between OB and LEC in neonatal mice. **(A)** Long-range projections of tdTomato fluorescently labeled OB M/T cells (left) when superimposed on a bright-field image showing the ventral (middle) and lateral (right) view of the whole brain of a P10 Tbet-cre;tdTomato mouse. **(B)** Unprocessed (left) and cleared (right) brain of a P10 mouse. **(C)** Cleared 500 μm-thick coronal section containing the OB of a Tbet-cre;tdTomato mouse showing MTCs (red) when counterstained with the nuclear marker DRAQ5 (blue). Inset, tdTomato-stained MTCs displayed at larger magnification. **(D)** MTC axons targeting LEC in a cleared 1 mm-thick coronal brain slice. Inset, axons of tdTomato-expressing MTCs when counter-stained with DRAQ5 (blue) and displayed at larger magnification. **(E)** Photographs of a 100 μm-thick coronal section from a P8 mouse depicting retrogradely labeled neurons in the OB (right) after injection of FG into the LEC (left) at P3. Inset, FG-labeled MTCs displayed at larger magnification. **(F)** Photographs of a 50 μm-thick coronal section from a P8 mouse depicting retrogradely labeled neurons in LEC (right top) after injection of FG into OB (100 μm-thick coronal section, left) at P4 (LEC, lateral entorhinal cortex; PIR, piriform cortex; PMCo, posteromedial cortical amygdaloid nucleus; rf, rhinal fissure; af, amygdaloid fissure). Counterstaining for GABA (red) shows no overlap between GABA positive (red) and FG (white) labeled cell bodies (right bottom).

Second, we assessed the spatial organization of feedback projections from LEC to OB. Unilateral injection of FG confined to OB of P3-4 mice (n=12) led to bright fluorescent back-labeling of parental cell bodies in ipsilateral LEC that project to OB of P8-10 mice (Fig 1F). Their density was lower when compared to the cells detected in ipsilateral PIR. Most labeled neurons were located in layer II and III (88.40%, 259/293, 3 pups, 11 sections). To examine the neurochemical identity of entorhinal neurons projecting to OB, we counter-stained the FG-labeled neurons for GABA and CamKII. While most OB-projecting neurons (99.66%, 292/293) were negative for GABA, hence glutamatergic, a small fraction 0.34%, 1/293) was GABA-positive. Similarly, CamKII staining revealed that the large majority, but not all FG-labeled cells, were glutamatergic (data not shown). These data indicate that top-down projections from LEC to OB can be either excitatory or inhibitory, as recently described for adult mice (33).

Taken together, the results of morphological investigation show that afferent and efferent projections couple neonatal LEC and OB. While glutamatergic MTCs axons target entorhinal layer I, glutamatergic and few GABAergic neurons in superficial layers of LEC innervate the OB.

### Continuous respiration-related activity and discontinuous theta bursts entrain the neonatal OB

Despite abundant data on morphological development, the functional maturation of OB is still largely unknown. In contrast to the retina and cochlea, which lack stimulus sensitivity at early stages of postnatal development and only generate spontaneous activity, the OB processes olfactory input already at birth (34). To elucidate the patterns of activity in the neonatal OB, we performed multi-site extracellular recordings of local field potential (LFP) and multiple unit activity (MUA) from the mitral cell layer in the dorsal and ventral OB of P8-10 mice *in vivo* (n=49). Signal reversal between the internal plexiform layer (IPL) and external plexiform layer (EPL), as well as the large MTC spikes served as physiological markers for confirming the position of recording electrode set according to stereotaxic coordinates. In addition, the location of DiI-labeled electrodes was confirmed after histological investigation *post mortem* (Fig 2A, S1 Fig A, B).

**Fig 2.**
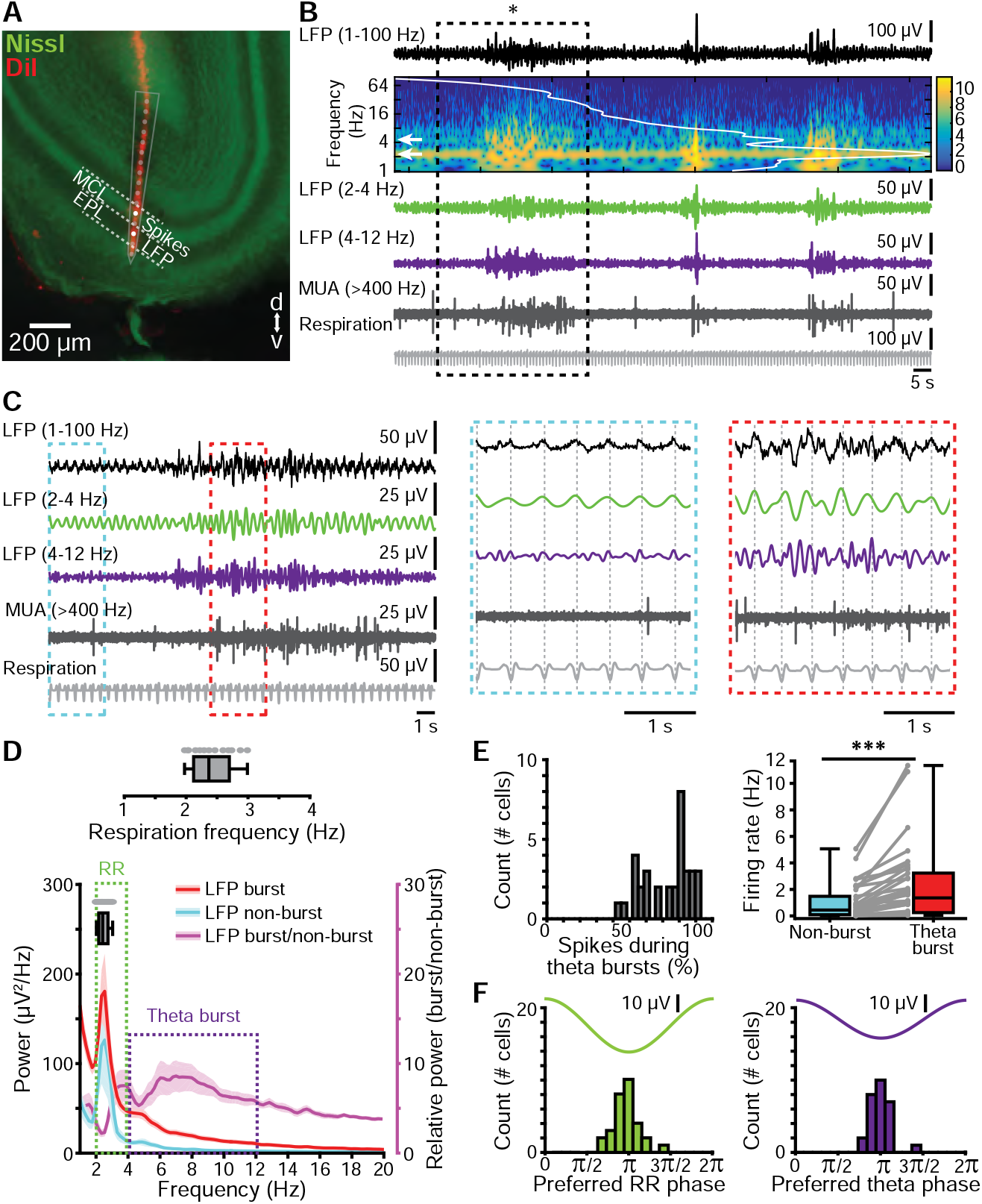
Continuous and discontinuous patterns of oscillatory activity in the neonatal olfactory bulb. **(A)** Digital photomontage reconstructing the track of the multi-site DiI-labeled recording electrode (red) in a Nissl-stained (green) 100 μm-thick coronal section including the OB from a P9 mouse. The dots (gray) show the position of the 16 recording sites of the silicon probe and the recording channels (white) in the mitral cell layer (MCL) and external plexiform layer (EPL) that were used for spike and LFP analysis, respectively. **(B)** LFP recording of the oscillatory activity in the OB of a P10 mouse displayed band-pass filtered in different frequency bands and accompanied by the wavelet spectrogram (white line represents time-averaged power of the trace; white arrows point towards peak frequency values) as well as simultaneously recorded MUA (high-pass filter >400 Hz) and respiration. **(C)** Characteristic slow continuous oscillatory activity and theta bursts from the trace shown in b when displayed at higher magnification. Insets, continuous oscillatory activity in relationship with respiration (left, blue) and a discontinuous theta burst (right, red). **(D)** Power spectra (mean ± SEM) of LFP in OB during non-burst activity (blue) and discontinuous bursts (red) as well as of theta bursts normalized to non-bursting activity (purple). The respiration frequency was depicted as horizontal bar and expanded at larger scale (top). **(E)** Temporal relationship between neuronal firing and network oscillations in OB. Left, histogram showing the percentage of spikes occurring during theta burst for all clustered units. Right, box plot depicting the firing rates of OB units during non-burst periods and theta burst periods. Gray dots and lines correspond to individual cells (Wilcoxon signed-rank test, ***p < 0.001). **(F)** Histograms depicting the phase locking of OB cells to RR (left) and theta activity (right). Only significantly locked cells were used for analysis.

Two patterns of coordinated activity were detected in OB (Fig 2B, C). First, we recorded continuous low amplitude oscillations with slow frequency peaking at 2-4 Hz. Given their temporal correlation and frequency overlap with respiration (median frequency: 2.37 Hz, iqr: 2.12-2.70 of chest movements) (Fig 1C, D), we defined this activity as respiration-related rhythm (RR). The RR reversed at the level of MTC layer and had larger amplitudes in EPL and glomerular layer when compared to the activity in MTC layer (data not shown). Its temporal relationship to the phase of the respiratory cycle differed between layers; the peak of RR cycle in the granule cell layer (GCL) and its trough in EPL and glomerular layer correlated with exhalation. Second, we recorded discontinuous high amplitude oscillatory events with spindle shape in the neonatal OB (Fig 2B, C). These events had faster frequencies when compared to RR with a peak within theta frequency band (4-12 Hz) (Fig 2D). Given their resemblance in shape and frequency dynamics to previously characterized oscillatory events in neonatal cortical areas (9-11, 35), these events were classified as theta bursts.

As reported for adult OB, prominent spiking characterized neonatal MTCs. Analysis of single unit activity (SUA) after principal component analysis (PCA)-based sorting of units revealed that the majority (80%) of spikes occurred during theta bursts. The firing rate during bursts (median: 1.36 Hz, iqr: 0.25-3.23 Hz) was significantly (p=3.65×10^−7^, Wilcoxon signed-rank test, n=34 cells from 14 animals) augmented when compared to non-bursting periods (median: 0.44 Hz, iqr: 0.09-1.48 Hz) (Fig 2E). To assess the temporal relationship between oscillatory OB rhythms and MTC firing, we estimated the coupling strength between SUA and RR as well as between SUA and theta bursts by calculating the pairwise phase consistency (PPC), a bias-free measure of rhythmic neuronal synchronization (36). Both rhythms similarly timed MTC firing (RR: median PPC: 0.21, iqr: 0.20-0.22 vs. theta burst: median PPC: 0.21, iqr: 0.20-0.21., p=0.1664, Wilcoxon signed-rank test, 2 outliers removed, n=32 cells, Fig 2F).

In adults, dorsal and ventral OB subdivisions have distinct physiology and function. MTC axons that originate in the dorsal OB are known to strongly project to amygdala and mediate innate odor responses, whereas ventral OB accounts for processing of learned odorants (37, 38). To assess whether distinct activity patterns entrain the dorsal vs. ventral OB at neonatal age, we compared RR and theta bursts from both subdivisions (S1 Fig). The power of RR was similar in both sub-divisions (dorsal: median 233.12 μV^2^, iqr: 153.42-418.09, n=7; ventral: median 335.75 μV^2^, iqr: 195.91-452.52, n=10; p=0.54, Wilcoxon rank-sum test). Similarly, theta burst occurrence (dorsal: median 4.65 bursts/min, iqr: 3.87-5.55; ventral: median 5.09 bursts/min, iqr: 4.27-5.30, p=0.74, Wilcoxon rank-sum test), duration (dorsal: median 6.76 s, iqr: 4.44-8.86 s; ventral: median 3.54 s, iqr: 1.65-5.01 s; p=0.09, Wilcoxon rank-sum test), amplitude (dorsal: median 73.80 μV, iqr: 59.56-75.37; ventral: median 66.59 μV, iqr: 59.10-72.37; p=0.67, Wilcoxon rank-sum test) and relative power (dorsal: median 493.98 Hz, iqr: 430.71-763.00; ventral: median 452.63 Hz, iqr: 395.3-1071.6, p=0.96, Wilcoxon rank-sum test) were comparable across OB subdivisions. These data indicate that the dorsal and ventral OB show similar activity at early postnatal age. Except otherwise indicated, further investigation focused on the ventral OB subdivision, taking into account its role for learning processes in relation with the limbic system (38).

Coordinated patterns in the sensory periphery have been reported to critically depend on the brain state, diminishing or even disappearing in the presence of anesthetics (39, 40). In contrast, early oscillations in the developing brain have often been investigated in the presence of urethane anesthesia (9, 35, 41, 42). Rodent pups spend most of the time sleeping. The sleep-mimicking action of urethane might explain the similar patterns of neuronal activity previously observed in anesthetized and sleeping rodent pups (14, 43). To assess the urethane influence on RR and theta bursts, we recorded from both ventral (n=12) and dorsal OB (n=6) of neonatal mice before and after urethane injection. Anesthesia did not change the overall structure of OB activity, with continuous RR and discontinuous theta bursts persisting (S2 Fig A, S1 Table). Both the power of RR and the occurrence of theta bursts remained unchanged (S2B Fig). However, urethane anesthesia profoundly reduced theta burst duration (S2B Fig), augmenting those time windows lacking theta band activity and therefore, the fragmented appearance of neonatal activity in OB (S2B Fig).

These data indicate that, independent of OB subdivision and brain state, the neonatal OB shows two main patterns of early oscillatory activity, continuous RR activity and discontinuous theta bursts.

### Mechanisms underlying the generation of continuous and discontinuous oscillatory activity in the neonatal OB

To elucidate the mechanisms contributing to the generation of continuous RR and discontinuous theta bursts in the OB of neonatal mice, we used two experimental approaches. First, the temporal coupling between respiration and continuous 2-4 Hz oscillations in OB suggests that nasal air flow contributes to RR generation. To test this hypothesis, we reduced the nasal air flow by unilateral naris occlusion in P8-10 pups (n=12) using a previously developed protocol (44, 45). MUA and oscillatory activity of OB were recorded before and after naris occlusion with silicon adhesive (data not shown). While unilateral deprivation did not change the overall structure of OB activity patterns, it reduced the RR power from 396.05 μV^2^ to 293.30 μV^2^ (baseline: iqr 232.58-570.88 μV^2^; occlusion: iqr 136.10-410.14 μV^2^, p=0.0009, Wilcoxon signed-rank test). By contrast, the theta bursts in OB were not affected by naris occlusion (baseline: median: 643.45 Hz, iqr: 342.6-1009.7; occlusion: median 700.35 Hz, iqr 284.5-1240.8; p=0.91, Wilcoxon signed-rank test). Correspondingly, the firing rate during RR (baseline: median 1.18 Hz, iqr 0.26-2.45) as well as coupling strength (i.e. PPC) between units and RR (baseline: median 8.50×10^−4^, iqr 0-0.0084, one outlier removed) decreased after naris occlusion (firing rate: occlusion: median 0.75 Hz, iqr 0.25-1.91, p=0.021. Wilcoxon signed-rank test; coupling strength: occlusion: median −3.09×10^−5^, iqr −2.76^×104^-1.56×10^−4^; p=0.049, Wilcoxon signed-rank test, one outlier removed). The temporal structure (coupling strength for baseline: median 2.09×10^−4^, iqr: – 0.0001-0.0015; occlusion: median −1.19×10^−4^, iqr −3.54×10^−4^-1.59×10^−4^; p=0.19, Wilcoxon signed-rank test, one outlier removed) of OB firing in relationship with theta bursts remained unchanged after naris occlusion. Thus, RR activity, but not theta bursts critically depends on nasal air flow.

The second experimental approach aimed at assessing the role of MTCs, the OB projection neurons, to the generation of coordinated patterns of oscillatory activity. For this, we selectively manipulated MTC firing by light in P8-10 pups bred from crossing hemizygous Tbet-cre mice with R26-homozygous R26-ArchT-EGFP mice. By these means, MTCs of cre-positive mice selectively expressed the proton pump ArchT fused with EGFP. Already at P8, the fusion protein expression was robust both in MTC somata (S3A Fig) and axonal projections targeting LEC, PIR and posterior cortical amygdala (Fig 3A). Cre-negative mice were used as controls.

**Fig 3.**
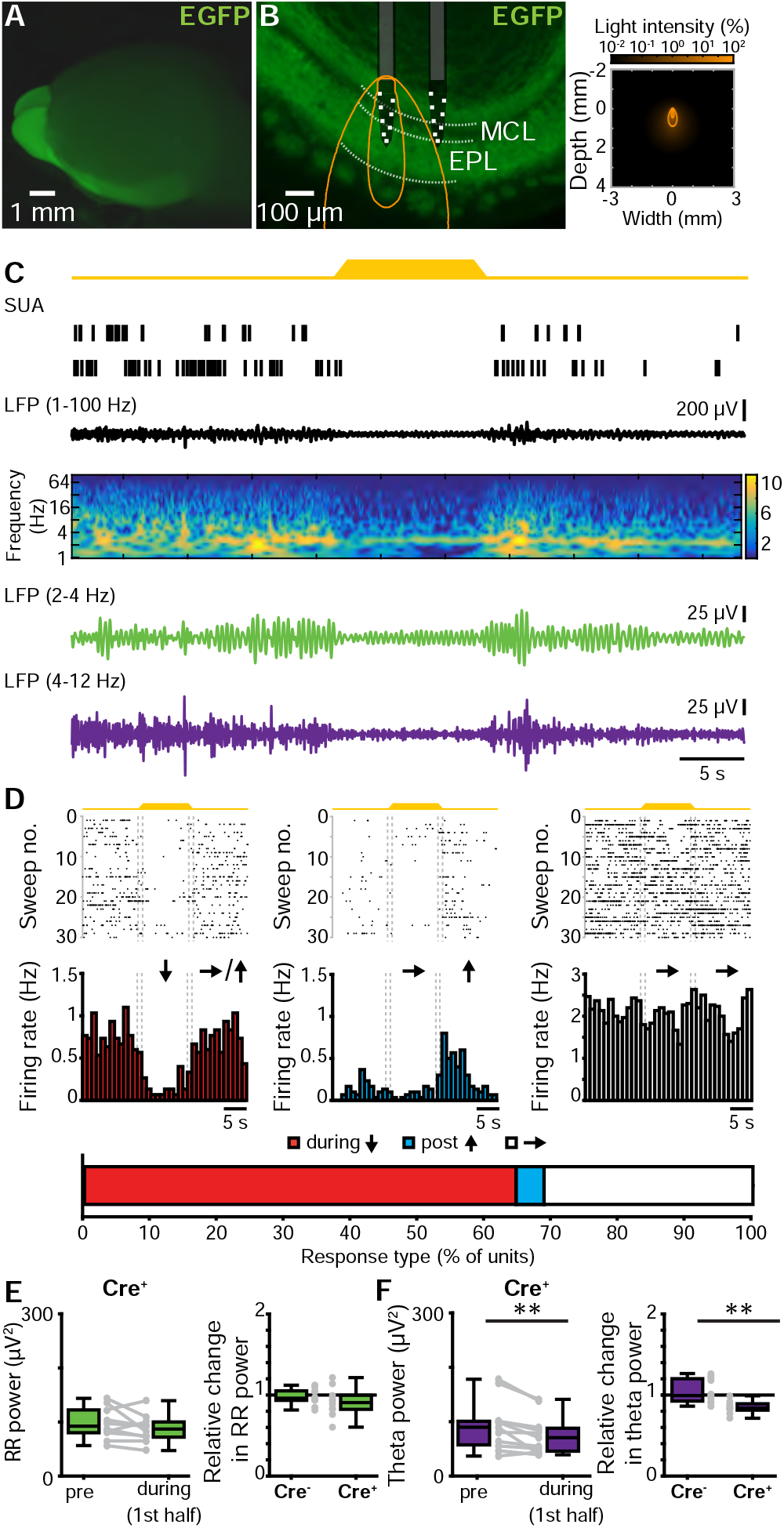
Effects of optogenetic silencing of MTCs on the patterns of oscillatory activity in the neonatal OB. (**A**) Photograph of the brain of a P8 cre-positive Tbet-cre;ArchT-EGFP mouse (left) showing EGFP-fluorescent MTCs cell bodies and their projections. **(B)** Left, photograph of a 100 μm-thick coronal section including the OB from a P8 cre-positive Tbet-cre;ArchT-EGFP mouse. The position of recording sites in MCL and EPL layers is marked by white squares. The light guide ending just above the recording sites is shown in gray. The iso-contour lines of light spreading calculated using Monte Carlo simulation are shown in yellow. Right, propagation of light intensity in the brain as predicted by Monte Carlo simulation. Yellow lines correspond to the iso-contour lines for light power of 1 and 10 mW/mm^2^, respectively. **(C)** Neuronal firing (SUA) and LFP band-pass filtered for different frequency bands (broad 1-100 Hz, RR 2-4 Hz, theta 4-12 Hz) in response to light (yellow, 594 nm) stimulation of MTCs in a P8 cre-positive Tbet-cre;ArchT-EGFP mouse. Traces are accompanied by the color-coded wavelet spectrogram of LFP shown at identical timescale. **(D)** Raster plots and peri-stimulus time histograms displaying the firing of MTCs in response to light stimulation. The color-coded bar (bottom) displays the fraction of cells that responded with a firing decrease during stimulus (red), constant firing during stimulus but a firing increase post-stimulus (blue) and unchanged firing rate (white). **(E)** Box plots displaying the absolute power before and during light stimulation in cre-positive pups (left) and the relative change of RR activity in neonatal OB of cre-positive and cre-negative mice (right). Gray dots and lines correspond to individual animals. **(F)** Same as E for discontinuous theta bursts (**p < 0.01, left: signed-rank test, right: rank-sum test).

In a first experiment, we tested the efficiency of light-dependent MTC silencing in neonatal OB by performing whole-cell patch-clamp recordings from biocytin-filled EGFP-positive neurons (n=7 cells) in coronal slices containing the OB of P8-10 R26-heterozygous Tbet-cre;R26-ArchT-EGFP mice (n=5) (S3A Fig). Yellow light pulses (595 nm, 5 s, 0.2-0.6 mW) triggered MTC hyperpolarization from −49.96 mV to −58.39 mV (baseline: iqr −57.28-45.61 mV; light administration: iqr −63.10-48.89 mV, Wilcoxon signed-rank test, p=0.0078) and, consequently, inhibition of firing (S3B Fig). Since MTCs are strongly interconnected within local circuits, we tested whether light pulses caused MTC silencing also in the presence of synaptic inputs. To mimic such inputs, we paired the light stimulation with depolarizing current pulses of different intensities. Upon injections ≤ 60 pA, light stimulation still efficiently blocked action potential discharge in ArchT-EGFP-expressing MTCs (S3C Fig).

Next, we assessed the contribution of MTC firing to the patterns of oscillatory activity in OB by performing extracellular recordings of LFP and MUA in OB of P8-10 R26-heterozygous cre-positive (n=12) and cre-negative (n=11) Tbet-cre;ArchT-EGFP mice. Upon *in vivo* light stimulation (Fig 3A), the majority (64.58%, 31/48) of MTCs responded with a pronounced firing rate decrease from a median of 1.2 Hz (iqr 0.66-2.26) before to 0.45 Hz (iqr 0.13-0.99) during light exposure. None of the units augmented the firing during illumination and only few units (4.14%, 2/48) showed a post-stimulus firing increase (Fig 3D). Some units (31.25%, 15/48), most likely non-MTCs located close to the mitral cell layer, did not respond to light stimulation. Local silencing of MTCs modified the coordinated activity of OB. The properties of RR and theta bursts (theta burst power: p=0.23, Wilcoxon rank-sum test) were largely similar in cre-negative and cre-positive mice under control conditions (i.e. no light stimulation). Only the power of RR activity was slightly different (RR power: p=0.03, Wilcoxon rank-sum test). Upon light stimulation the RR power in cre-positive pups did not change (pre stimulus: median 92.27 μV^2^, iqr 80.12-122.36; during stimulus: median 86.99 μV^2^, iqr 71.96-100.06, p=0.2324, Wilcoxon signed-rank test, one outlier removed). In contrast, theta power in cre-positive pups significantly decreased during light stimulation (pre stimulus: median 89.73 μV^2^, iqr 57.28-100.73; during stimulus: median 70.58 μV^2^, iqr 45.70-87.91, p=0.0049, Wilcoxon signed-rank test, one outlier removed). The theta responses to light differed between cre-positive (median 0.84 μV^2^, iqr 0.81-0.89) and cre-negative pups (median 0.99 μV^2^, iqr 0.93-1.20, p=0.0024, Wilcoxon rank-sum test, 3 outliers from expression group removed), whereas for RR during light stimulus was similar in the two groups (cre-positive, median 0.91, iqr 0.82-1.0; cre-negative pups median 0.96, iqr 0.93-1.05, p= 0.3447, Wilcoxon rank-sum test, 1 outlier from control group, 2 outliers from expression group removed) (Fig 3E, F).

These data show that RR and theta bursts in the neonatal OB have different origin. While RR critically depends on nasal air flow, MTC activity is necessary for the entrainment of OB in theta bursts.

### Theta bursts in OB drive discontinuous oscillations and time the firing in the neonatal LEC

The presence of both direct axonal MTC-to-LEC projections and early patterns of oscillatory activity in OB led to the question of their relevance for the emergence of functional assemblies in the neonatal LEC. In contrast to the documented relevance of entorhinal output for developing limbic circuits (11), the role of sensory inputs for the functional maturation of LEC is still unknown.

Multi-site extracellular recordings of LFP and MUA from the layer II/III of LEC from P8-10 mice *in vivo* (n=11) (Fig 4A) confirmed the previously reported presence of discontinuous theta bursts with large amplitude (median 154.14 μV, iqr 101.10-191.65) and a duration of 5.15 s (iqr 4.13-8.48) (Fig 4B-D). They appear superimposed on a slow rhythm (2-4 Hz) that continuously entrains the neonatal LEC and has been overlooked in previous investigations. This slow pattern of activity that was present both during theta bursts (median area power 526.25 μV^2^, iqr 307.68-1171.85) and “silent” periods (median area power 86.57 μV^2^, iqr 52.55-344.43), temporally correlated with the simultaneously recorded respiration and was therefore, classified as entorhinal RR. These results demonstrate that the respiration-entrained brain rhythms, a powerful mechanism of long-range coupling (46), emerge early during development. Beside oscillatory patterns, neonatal LEC generates prominent firing concentrated during theta bursts (median 0.42 Hz, iqr 0.22-0.86 vs. nonbursting periods median 0.07 Hz, iqr 0.04 – 0.19, p=1.72×10^−10^, Wilcoxon signed-rank test, n=54 cells from 11 mice) (Fig 4E). We next assessed the coupling strength between firing and oscillatory activity. Similar fractions of entorhinal neurons were phase-locked to RR (75.93%, 41/54 units) and theta bursts (61.11%, 33/54 units, p=0.1, χ^2^(1)=2.7472). The strength of coupling assessed by PPC was also stronger for RR (median: 0.21, iqr: 0.20-0.22) and theta (median: 0.21, iqr: 0.20-0.21, p=2.98×10^−4^, Wilcoxon rank-sum test, 4 outliers removed, n=50 units), with most cells being locked to the trough of RR and theta oscillation (Fig 4F).

**Fig 4.**
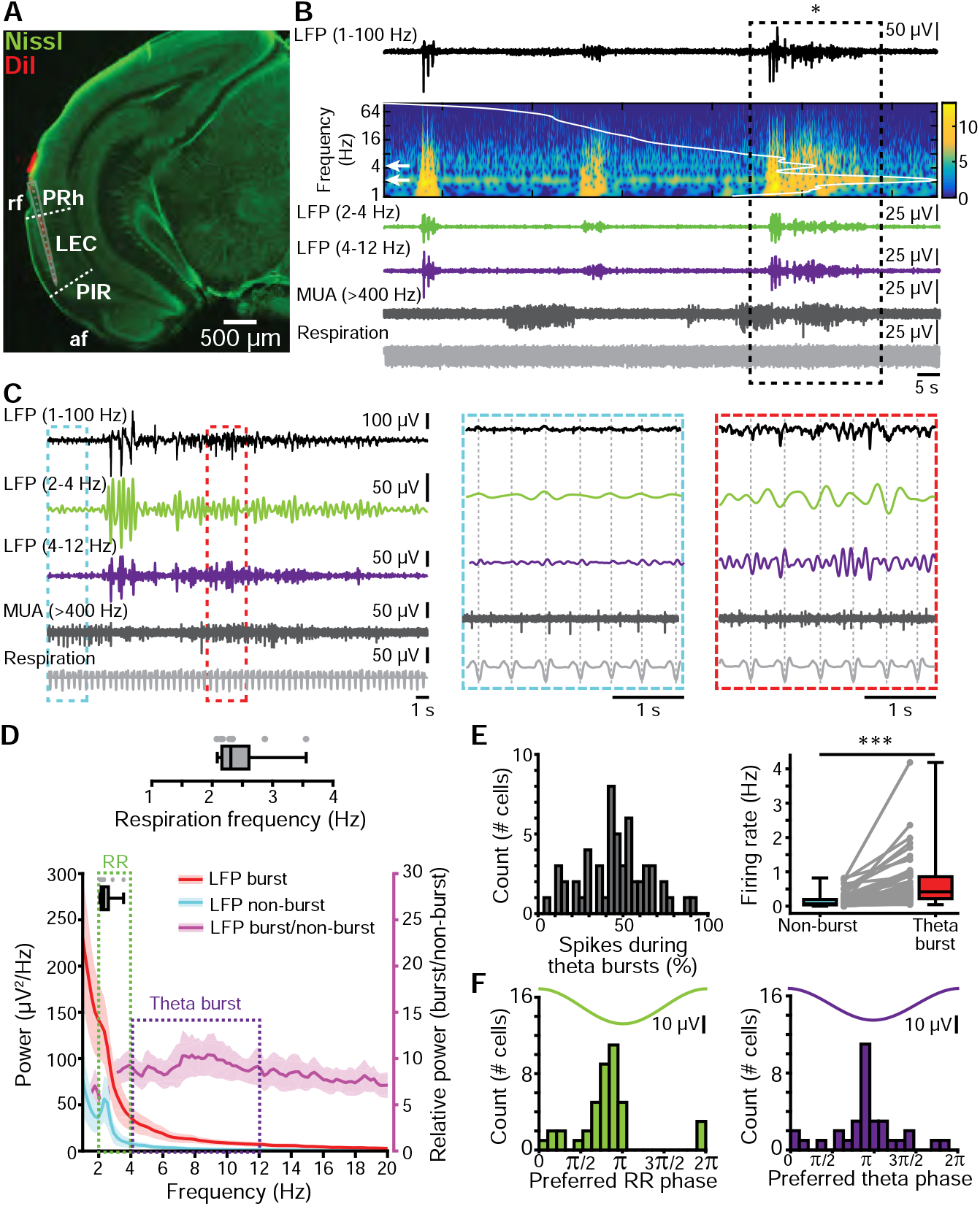
Continuous and discontinuous patterns of oscillatory activity in the neonatal LEC. **(A)** Digital photomontage reconstructing the track of the multi-site DiI-labeled recording electrode (red) in a Nissl-stained (green) 100 μm-thick coronal section including LEC from a P9 mouse. The gray dots show the position of the 16 recording sites. (PRh, perirhinal cortex; LEC, lateral entorhinal cortex; PIR, piriform cortex; rf, rhinal fissure; af, amygdaloid fissure). **(B)** LFP recording of the oscillatory activity in LEC of a P10 mouse displayed band-pass filtered in different frequency bands and accompanied by the wavelet spectrogram (white line represents time-averaged power of the trace) as well as simultaneously recorded MUA (high-pass filter >400 Hz) and respiration. **(C)** Characteristic slow continuous oscillatory activity and theta bursts from the trace shown in B when displayed at higher magnification. **(D)** Power spectra (mean ± SEM) of LFP in LEC during non-burst activity (blue) and discontinuous bursts (red) as well as of theta bursts normalized to non-bursting activity (purple). The respiration frequency was depicted as horizontal bar and expanded at larger scale (top). **(E)** Temporal relationship between neuronal firing and network oscillations in LEC. Left, histogram showing the percentage of spikes occurring during theta burst for all clustered units. Right, box plot depicting the firing rates of LEC units during non-burst periods and theta burst periods. Gray dots and lines correspond to individual cells (Wilcoxon signed-rank test, ***p < 0.001). **(F)** Histograms depicting the phase locking of LEC neurons to RR (left) and theta activity (right). Only significantly locked cells were used for analysis.

Simultaneous recordings from OB and LEC (n=9) of neonatal mice gave first insights into their dynamic coupling (Fig 5A). While both areas showed similar oscillatory activity, their power significantly differed. Both RR power (OB: median 143.62 μV^2^, iqr 78.03-247.78; LEC: median 109.91, iqr: 26.72-110.51, p=0.0499, Wilcoxon signed-rank test, 1 outlier removed) and theta power (OB: median 193.39 μV^2^, iqr 97.47-262.01, LEC: median 112.37 μV^2^, iqr 43.38-127.36, p= 0.0273) were higher in OB as compared to LEC (Fig 5B). Analysis of the temporal correspondence of theta bursts in OB and LEC revealed that 48.70% of them cooccurred with more than 60% temporal overlap. The coupling strength assessed by imaginary spectral coherence, which excludes synchrony effects due to volume conductance (47), revealed that the OB-LEC coupling is evident in both slow frequencies (i.e RR) and theta band (i.e. theta bursts) (Fig 5C). In line with anatomical data, we detected no differences in the coupling of dorsal and ventral OB with LEC. Both relative occurrence of cooccurring events (dorsal: median 27.04 %, iqr 20.08-33.98 %; ventral: median 21.83 %, iqr 15.66-35.14 %; p=0.67, Wilcoxon rank-sum test) and mean imaginary coherence in both RR (dorsal: median 0.11 Hz, iqr 0.09-0.14 Hz; ventral: median 0.08 Hz, iqr 0.05-0.10 Hz; p=0.13, Wilcoxon rank-sum test) and theta frequency range (dorsal: median 0.07 Hz, iqr 0.06-0.11 Hz; ventral: median 0.06 Hz, iqr 0.06-0.09 Hz; p=0.54, Wilcoxon rank-sum test) were similar for dorsal and ventral OB in relationship to LEC (S1 Fig). These data are in line with anatomical investigations in adult mice (48) as well as with our tracing data (Fig 1), showing that, FG injections into neonatal LEC leads to homogenous MTC labeling throughout the OB.

**Fig 5.**
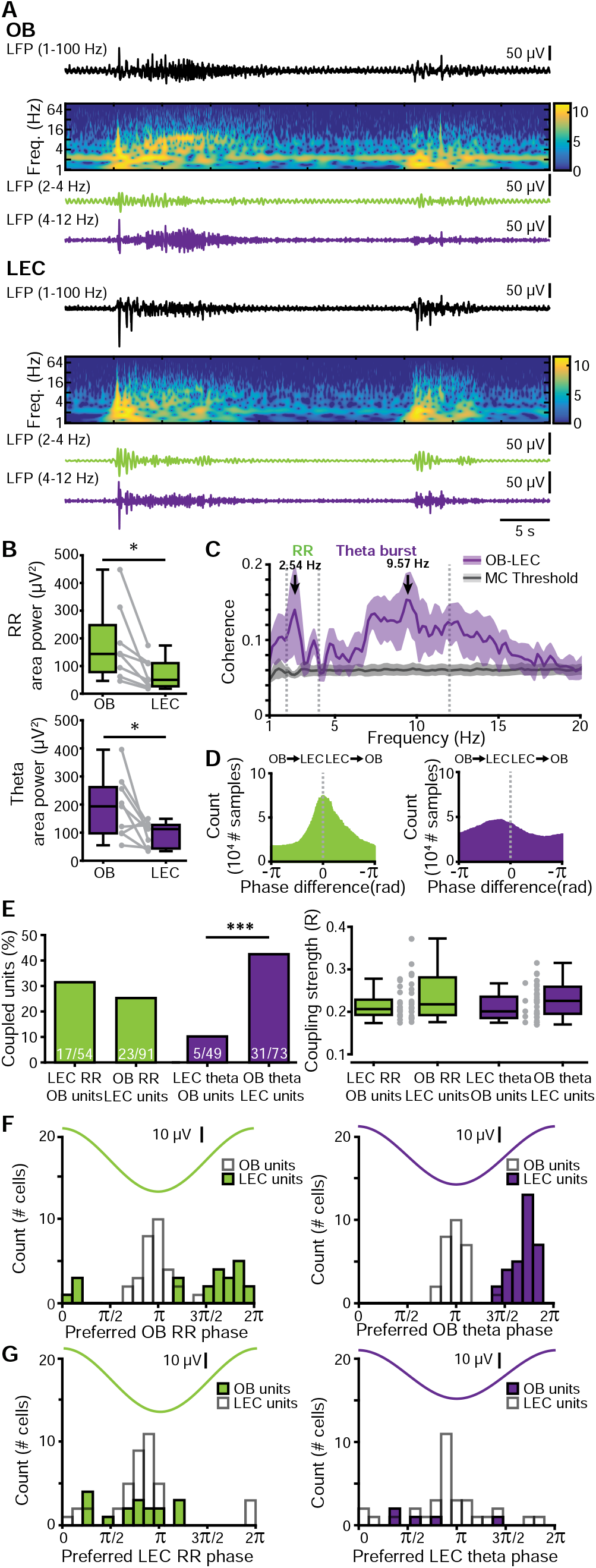
Frequency-dependent functional coupling between neonatal OB and LEC. **(A)** Characteristic traces of band-pass filtered LFP recorded simultaneously in OB (top) and LEC (bottom) of a P9 mouse, displayed together with wavelet spectrograms showing the frequency content. Note the temporal correlation between discontinuous theta bursts in both areas. **(B)** Boxplots displaying RR power (top, green) and theta burst power (bottom, purple) in OB and LEC. Gray lines and dots correspond to individual pups. (*p<0.05, Wilcoxon signed-rank test). **(C)** Plot of imaginary part of coherence between OB and LEC showing prominent peaks in RR and theta band. The gray line corresponds to the significance threshold as assessed by Monte Carlo simulation. **(D)** Histograms of phase differences between RR (left, green) and theta (right, purple) activity recorded simultaneously in OB and LEC. **(E)** Left, bar diagram displaying the percentage of OB units coupled to the RR (green) and theta bursts (purple) in LEC and the percentage of LEC units coupled to the RR (green) and theta bursts (purple) in OB. Right, box plot showing the coupling strength of OB cells significantly locked to LEC oscillations (green: RR, purple: theta bursts) and of LEC cells significantly locked to OB oscillations (green: RR, purple: theta bursts). Gray dots correspond to individual cells (χ^2^ test of proportions ***p < 0.001). **(F)** Histograms showing the distribution of preferred phases of LEC cells significantly locked to RR (left) and OB theta bursts (right) in neonatal OB. For comparison, histograms of OB cells locked to the respective OB rhythm are plotted as white bars. **(G)** Histograms showing the distribution of preferred phases of OB cells significantly locked to RR (left) and theta bursts (right) in neonatal LEC. For comparison, histograms of LEC cells locked to the respective LEC rhythm are plotted as white bars.

To assess the influence of anesthesia on entorhinal activity patterns and coupling between LEC and OB, we recorded both areas in mouse pups before and after urethane i.p. injection (n=18). Urethane did not change the overall spectral distribution of activity patterns in LEC. As in the non-anesthetized state, RR and theta bursts were the main patterns of entorhinal activity, yet the RR power decreased and theta power augmented under urethane action (S4A Fig, S1 Table). Urethane affected the duration of theta bursts and slightly increased their occurrence (S4B Fig). The synchrony between OB and LEC varied in magnitude, but not frequency distribution. The imaginary coherence peaked at 2-4 Hz and at 5 to 20 Hz, corresponding to RR and theta-beta frequencies, respectively. While mean RR coherence did not differ between states (p=0.17, Wilcoxon rank-sum test, 2 outliers removed), theta coherence was higher in the presence of urethane (p=0.0034).

These data indicate that, independent of brain state and anatomical subdivision, OB and LEC couple tightly, both being synchronized in continuous RR and discontinuous theta oscillations at neonatal age.

Since feed-forward projections from MTCs to LEC are dense, whereas feed-back projections from LEC to OB are rather sparse, we asked whether the functional coupling between the two areas is directed, and if so, whether directionality is frequency-specific. To estimate the directionality of OB-LEC coupling, we used two approaches. First, we assessed the phase lag between LFP in OB and LEC. While the phase lag for continuous RR was centered to 0, it peaked in negative range for theta bursts, indicating that OB theta bursts most likely drive LEC theta oscillations (Fig 5D). Second, we analyzed the temporal relationship between spiking activity in one area and either LFP or spiking in the other area. For RR, a similar number of clustered units in OB and LEC were phase-locked to RR in LEC (31.48%, 17/54) and OB (25.27%, 23/91 p=0.54, χ^2^(1)=0.38, χ^2^ test of proportions), respectively and their coupling strengths were comparable (p=0.35, Wilcoxon rank-sum test, OB cells to LEC RR: median 0.21, iqr 0.19-0.23; LEC cells to OB RR: median 0.22, iqr 0.19-0. 28, Fig 5E). In contrast, a significantly higher fraction of LEC neurons were phase-locked to theta bursts in OB (42.47%, 31/73) when compared to OB neurons timed by entorhinal theta phase (10.20%, 5/49, p=1.28×10-4, χ^2^(1)=14.67, χ^2^ test of proportions, Fig 5E). The coupling strengths of these neuronal populations, however, were comparable (p=0.44, Wilcoxon rank-sum test, OB cells to LEC theta: median: 0.22, iqr: 0.19-0.24; LEC cells to OB theta: median: 0.23, iqr: 0.20-0.26, one outlier removed from OB theta-LEC units group). Notably, MTCs preferentially fire during the trough of RR activity and theta bursts in OB, whereas LEC cells preferentially fire on the rising phase after the trough of OB rhythms, indicating that MTC firing precedes LEC cell firing by about a third of a cycle (Fig 5F, G).

Together, these data suggest that the continuous RR rhythm is not involved in directed information flow within OB-LEC circuits, whereas theta bursts in OB drive the oscillatory entrainment of LEC.

### Pharmacological blockade of OB firing diminishes the slow and fast oscillatory activity but not the coupling of OB-LEC circuits

To confirm the functional long-range coupling between OB and LEC, we pharmacologically abolished the neuronal activity by unilateral pressure-injection of the voltage-dependent sodium channel (hence action potential) blocker lidocaine (4% in sterile saline) into OB. Extracellular recordings of LFP and MUA were performed simultaneously from OB and LEC of mice (n=8) before and after lidocaine injection *in vivo* (Fig 6A). The injected lidocaine volume of 4 μl was proven to not spread across the borders of OB (Fig 6B). Lidocaine abolished OB firing within ten minutes of injection from a median baseline firing rate of 1.97 Hz (iqr: 0.77-2.80) to 0.00 Hz (iqr: 0.00-0.02). A partial recovery was observed after 30-40 min (χ^2^(7)=45.04, p=1.34×10^−7^, Friedman test, with Wilcoxon signed-rank post hoc test with Bonferroni correction) (Fig 6D). The firing of entorhinal neurons was also significantly reduced after lidocaine treatment in OB from a median baseline firing rate of 2.4 Hz (iqr 1.46-3.60) to 0.52 Hz (iqr 0.32-1.09) within the first 10 minutes after injection (χ^2^ (7)=135.50, p=4.45×10^−26^, Friedman test, with Wilcoxon signed-rank post-hoc test with Bonferroni correction). The decrease of firing rates in both areas was accompanied by changes of oscillatory network activity. In OB, the power of RR (baseline: median 91.05 μV^2^; iqr 70.66-224.40; lidocaine: median 8.88, iqr 3.90-20.16; p=0.0078, Wilcoxon signed-rank test) as well as the occurrence (baseline: median 4.76 bursts/min, iqr 3.58-5.93; lidocaine: 1.03 bursts/min, iqr 0.76-1.56, p=0.0234, Wilcoxon signed-rank test), duration (baseline: median 4.41s, iqr 3.78-4.77; lidocaine: median 2.35, iqr 1.89-2.88, p=0.0078, Wilcoxon signed-rank test) and power (baseline: median 202.07 μV^2^, iqr 163.62-261.95; lidocaine: median 33.19, iqr 24.14-109.00. p=0.0156, Wilcoxon signed-rank test) of theta bursts were reduced. In LEC, the power of RR (baseline: median 55.88 μV^2^, iqr 42.48-134.05; lidocaine: median 17.71, iqr 9.07-53.80, p=0.0391, Wilcoxon signed-rank test) as well as the duration (baseline: median 4.41s, iqr 3.74-4.58; lidocaine: median 3.24, iqr 3.14-3.56, p=0.0156, Wilcoxon signed-rank test, one outlier removed) and power (baseline: median 148.11 μV^2^, iqr 115.76-191.62; lidocaine: median 75.50, iqr 65.99-116.63, p=0.0313, Wilcoxon signed-rank test, 2 outliers removed) of theta bursts were decreased after blockade of OB firing. While the coordinated activity substantially diminished in OB and LEC, the coupling between both areas was not affected by lidocaine. Coherence neither changed in slow frequencies (baseline: median 0.08, iqr 0.05-0.11; lidocaine: median 0.14, iqr 0.08-0.24, p=0.25, Wilcoxon signed-rank test) nor in theta frequency band (baseline: median 0.11, iqr 0.07-0.16; lidocaine: median 0.14, iqr 0. 10-0.18, p=0.54, Wilcoxon signed-rank test) after blocking OB firing (Fig 6E).

**Fig 6.**
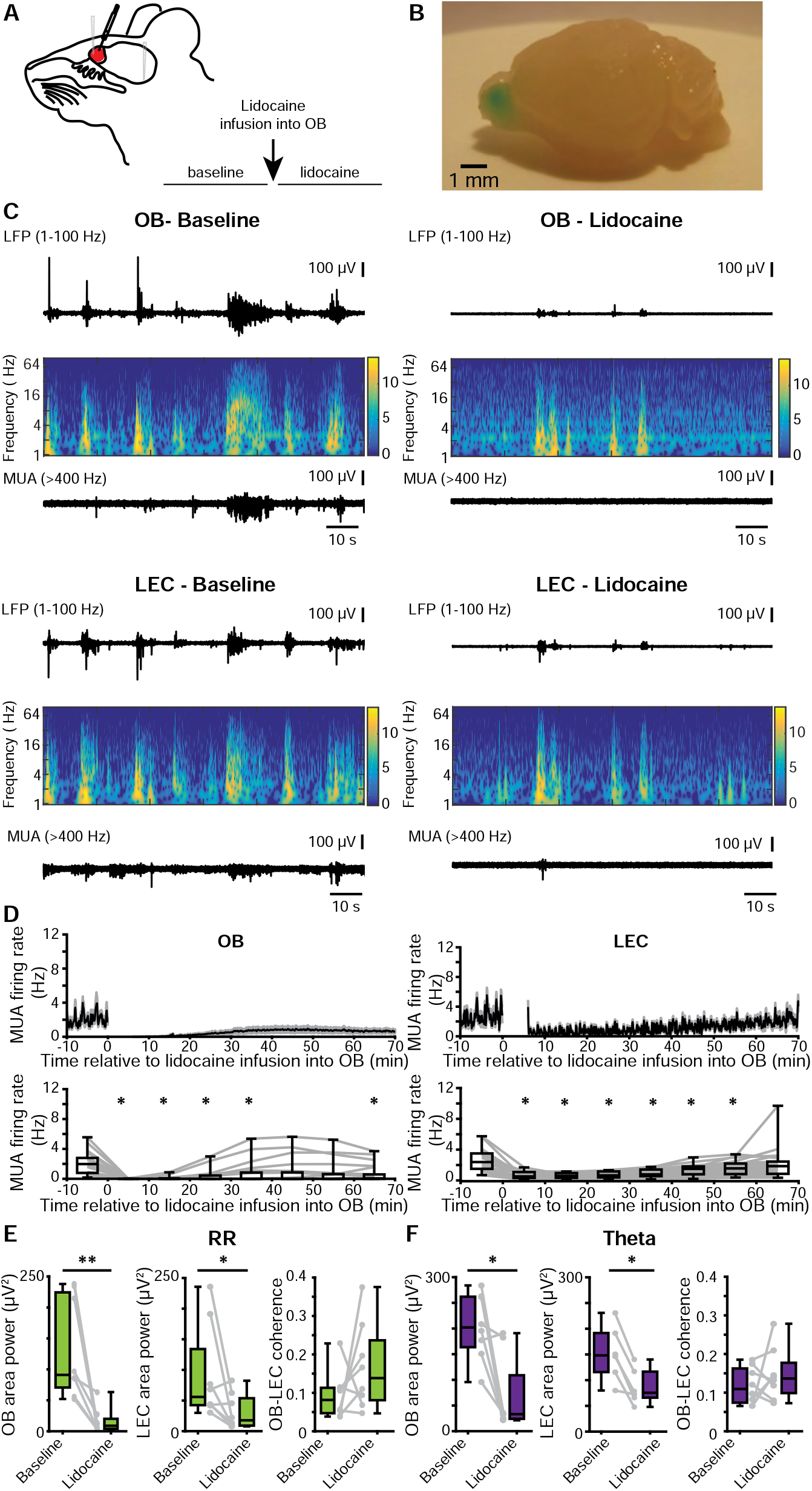
Effects of pharmacological blockade of neuronal firing in OB on patterns of oscillatory activity in OB-LEC circuits. **(A)** Schematic drawing of experimental protocol. **(B)** Photograph of the brain of a P10 mouse showing the confinement of injections to one hemisphere of the OB. For visualization, the same volume of methylene blue was used. **(C)** Characteristic LFP traces (black, filtered 1-100 Hz) recorded in OB (top) and LEC (bottom) of a P9 mouse before (left) and after (right) lidocaine infusion, displayed together with the wavelet spectrograms of the LFP and simultaneously recorded MUA. **(D)** Top, mean MUA firing rate in OB (left) and LEC (right) before and after lidocaine infusion. The time of infusion is considered 0. Bottom, box plots displaying the mean MUA in OB (left) and LEC (right) before and after lidocaine infusion (Friedmann test, Wilxocon signed-rank test with Bonferroni correction for post-hoc comparison, *p<0.0071). **(E)** Box plots displaying the power of RR activity in OB and LEC as well as the mean OB-LEC coherence in the RR band before and after lidocaine infusion. Gray dots and lines correspond to individual animals (Wilcoxon signed-rank test, *p < 0.05; **p < 0.01). **(F)** Same as E for the theta burst activity in neonatal OB and LEC.

These data indicate that blocking of neuronal firing in OB causes massive diminishment of coordinated activity in both OB and LEC but does not change the coupling strength between the two regions.

### Odors boost the oscillatory activity in neonatal OB and LEC and augment their fast frequency coupling

In contrast to other sensory systems that lack peripheral sensitivity for environmental stimuli during early postnatal development, the olfactory system is functional at birth. Therefore, the characterized coordinated patterns of oscillatory activity, RR and theta bursts, might have a dual origin, i.e. resulting from both spontaneous and/or stimulus-evoked activation of OB neurons. To gain first insights into the relevance of environmental stimuli on oscillatory activity and long-range entrainment of OB and LEC, we exposed P8-10 mouse pups to different odors. Prominent oscillatory discharge with slow and fast frequencies as well as MUA were induced in OB by olfactometer-controlled exposure to odors, such as octanal (10%) (Fig 7A). Intriguingly, we observed odor-evoked responses also in LEC, albeit at lower magnitude. Compared to theta bursts recorded in absence of stimuli (i.e baseline) and to responses to saline, these octanal-evoked responses had a higher amplitude of slow oscillations corresponding to RR both in OB (χ^2^ (2)= 36.05, p=1.49×10^−8^, Kruskal-Wallis test, Wilcoxon rank-sum test with Bonferroni correction as post-hoc test) and LEC (χ^2^ (2)= 13.80, p=0.001, Kruskal-Wallis test, Wilcoxon rank-sum test with Bonferroni correction as post-hoc test). Similarly, octanal augmented the amplitude of theta bursts in both regions (OB: χ^2^(2)= 36.30, p=1.31×10^−8^, Kruskal-Wallis test, Wilcoxon rank-sum test with Bonferroni correction as post-hoc test, LEC: χ^2^(2)= 20.52, p=0.000035, Kruskal-Wallis test, Wilcoxon rank-sum test with Bonferroni correction as post-hoc test) (Fig 7B, C, Table 1). In contrast to coordinated theta burst activity recorded in the absence of olfactory stimulation, evoked responses included beta band (15-30 Hz) activity. The amplitude of beta activity was significantly higher in the presence of octanal than during baseline or saline exposure both in OB (χ^2^(2)= 56.52, p=5.33×10^−13^, Kruskal-Wallis test, Wilcoxon rank-sum test with Bonferroni correction as post-hoc test) and LEC (χ^2^(2)= 31.94, p=1.16×10^−7^, Kruskal-Wallis test, Wilcoxon rank-sum test with Bonferroni correction as post-hoc test) (Fig 7B, C, Table 1). The presence of odor-driven OB activity confirms the maturity of receptor cells and odor-processing mechanisms in the olfactory system at early postnatal age. Moreover, the presence of odor-driven LEC activity indicates that coordinated activity from OB drives the oscillatory entrainment of LEC. To determine which oscillatory patterns are mainly involved in these directed OB-LEC interactions, we calculated the imaginary coherence between the two areas upon exposure to either saline or octanal. While the coherence in the slow frequency band (i.e. RR) was higher for odor-triggered events as compared to baseline events, it was similar for saline and octanal (χ^2^(2)= 23.22, p=9.06×10^−6^, Kruskal-Wallis test, Wilcoxon rank-sum test with Bonferroni correction as post-hoc test, Fig 7D, Table 1). In contrast, the coherence in fast frequencies significantly augmented in the presence of octanal when compared to saline-evoked or baseline events (theta: χ^2^(2)= 43.99, p=2.81×10^−10^, Kruskal-Wallis test, Wilcoxon rank-sum test with Bonferroni correction as post-hoc test, beta: χ^2^(2)= 48.48, p=2.98×10^−11^, Kruskal-Wallis test, Wilcoxon rank-sum test with Bonferroni correction as post-hoc test) (Fig 7D, Table 1). These data suggest that discontinuous bursts, either spontaneous or odor-induced, facilitate the long range OB-LEC coupling and boost local entrainment in beta band of entorhinal circuits.

**Fig 7.**
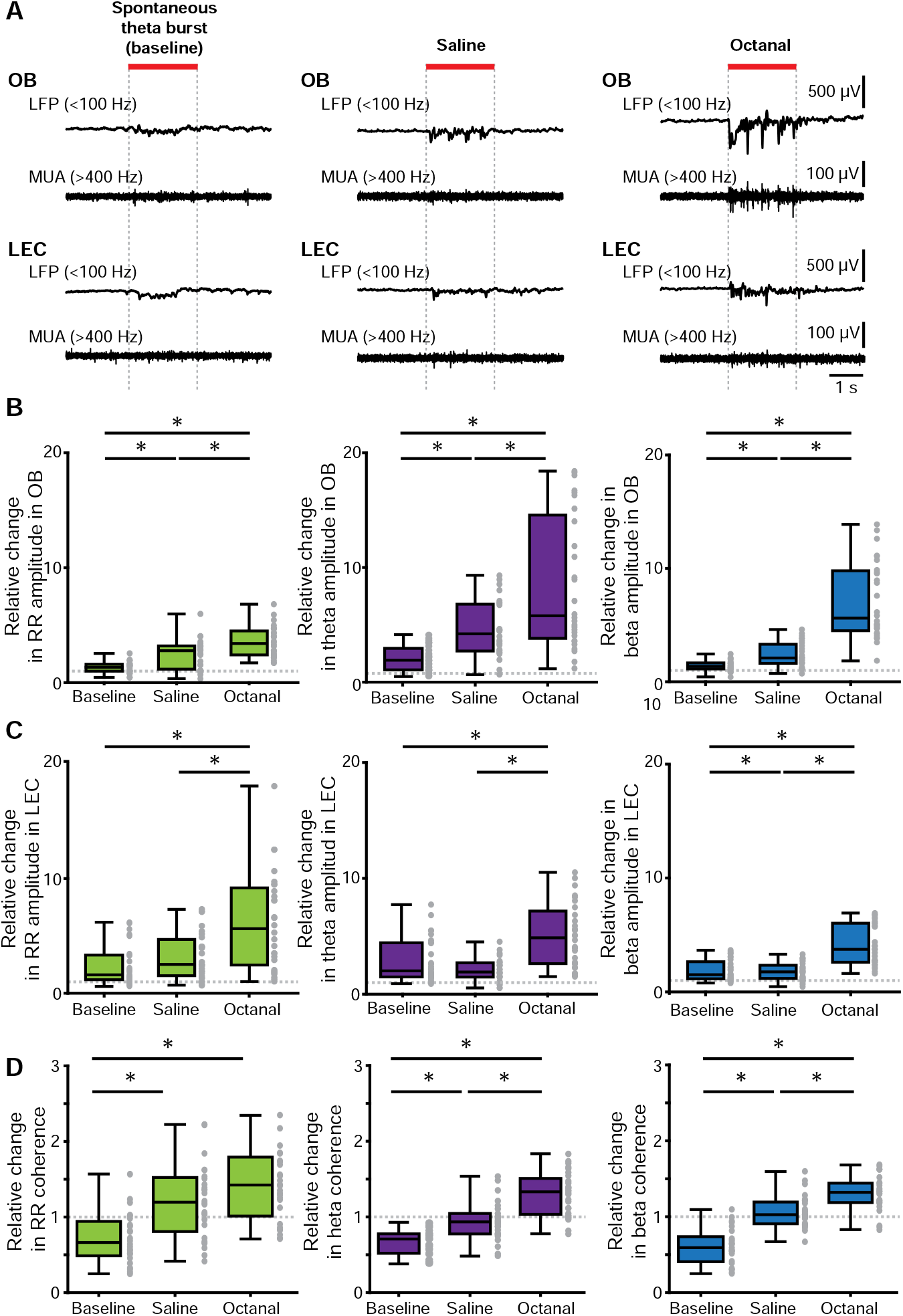
Odor-triggered activity patterns in OB of neonatal mouse. **(A)** Characteristic LFP traces (band-pass filtered 1-100 Hz) recorded in OB (top) and LEC (bottom) of a P9 mouse before (baseline, left) and after application of odors (saline, middle; octanal, right) displayed together with simultaneously recorded MUA. **(B)** Box plots showing odor-evoked changes in the amplitude of RR (left), theta (middle) and beta (right) activity in OB when normalized to baseline. **(C)** Same as B for LEC. **(D)** Odor-evoked relative changes in OB-LEC coherence in RR (left), theta (middle) and beta (right) band when normalized to baseline. Gray dots correspond to individual trials. (Kruskal-Wallis H test, Wilcoxon rank-sum test with Bonferroni correction as post-hoc test, *p<0.0167).

**Table 1.** (related to Figure 7). Quantification of odor-responses in neonatal OB-LEC networks.

|  | OB amplitude (relative change) |  |  |  | LEC amplitude (relative change) |  |  |  | OB-LEC coherence (relative change) |  |  |  |
| --- | --- | --- | --- | --- | --- | --- | --- | --- | --- | --- | --- | --- |
|  | No odor | Saline | Octanal | p | No odor | Saline | Octanal | p | No odor | Saline | Octanal | p |
| RR | 1.34<br>1.00-1.61 | 2.78<br>1.17 - 3.20 | 3.41<br>2.41 - 4.50 | <0.001 | 1.59<br>1.17-3.31 | 2.27<br>1.29 -4.45 | 5.38<br>2.21 - 8.94 | =0.001 | 0.69<br>0.50-1.0 | 1.18<br>0.76 - 1.49 | 1.38<br>0.90 - 1.73 | <0.001 |
| Theta | 1.90<br>1.05-2.93 | 4.22<br>2.72 - 6.80 | 5.79<br>3.82-14.57 | <0.001 | 2.01<br>1.48-4.43 | 1.19<br>1.48 - 2.71 | 4.87<br>2.64 - 7.18 | <0.001 | 0.71<br>0.55-0.79 | 0.93<br>0.78 - 1.05 | 1.33<br>1.03 - 1.51 | <0.001 |
| Beta | 1.37<br>1.14-1.68 | 2.10<br>1.62 - 3.31 | 5.61<br>4.50 - 9.79 | <0.001 | 1.52<br>1.15-2.66 | 1.76<br>1.18 - 2.34 | 3.74<br>2.60 - 6.03 | <0.001 | 0.61<br>0.42-0.76 | 1.03<br>0.91 - 1.19 | 1.32<br>1.19 - 1.44 | <0.001 |

The values are given as median and inter-quartile ranges and p-values (Kruskal-Wallis H test) for odor-evoked changes in oscillatory activity in RR, theta and beta band in OB and LEC, as well as OB-LEC coherence, when compared to baseline events are included.

## DISCUSSION

Assembling of neurons into functional networks during development is the pre-requisite for behavioral performance in adults. Entrainment of neurons into coordinated oscillatory rhythms represents a powerful assembling principle that has been initially identified to control the topographic organization of sensory systems (6, 8, 49, 50). More recently, patterns of coordinated activity have been characterized in developing limbic systems (9, 14, 15, 51, 52). However, it is still unclear whether sensory and limbic circuits adhere to similar assembling principles and how they interact during early development. In the present study, we tested the hypothesis that coordinated activity patterns of neuronal assemblies in neonatal OB contribute to the oscillatory entrainment of LEC, the gatekeeper of limbic circuits during development. Combining *in vivo* electrophysiology, optogenetics and pharmacology with anatomical tracing of projections, we demonstrate that (i) two major patterns of coordinated activity entrain the neonatal OB: continuous slow frequency oscillations temporally related to respiration and discontinuous theta band oscillations critically depending on MTC activity; (ii) both rhythms temporally couple the neonatal OB and LEC via dense direct axonal projections, with OB theta bursts boosting the oscillatory entrainment of entorhinal circuits; and (iii) olfactory stimuli augment oscillatory power, induce activity in fast frequency bands, and strengthen the coupling within OB-LEC circuits (Fig 8). These data reveal that endogenously-generated and stimulus-driven activities in OB control the oscillatory entrainment of LEC.

**Fig 8.**
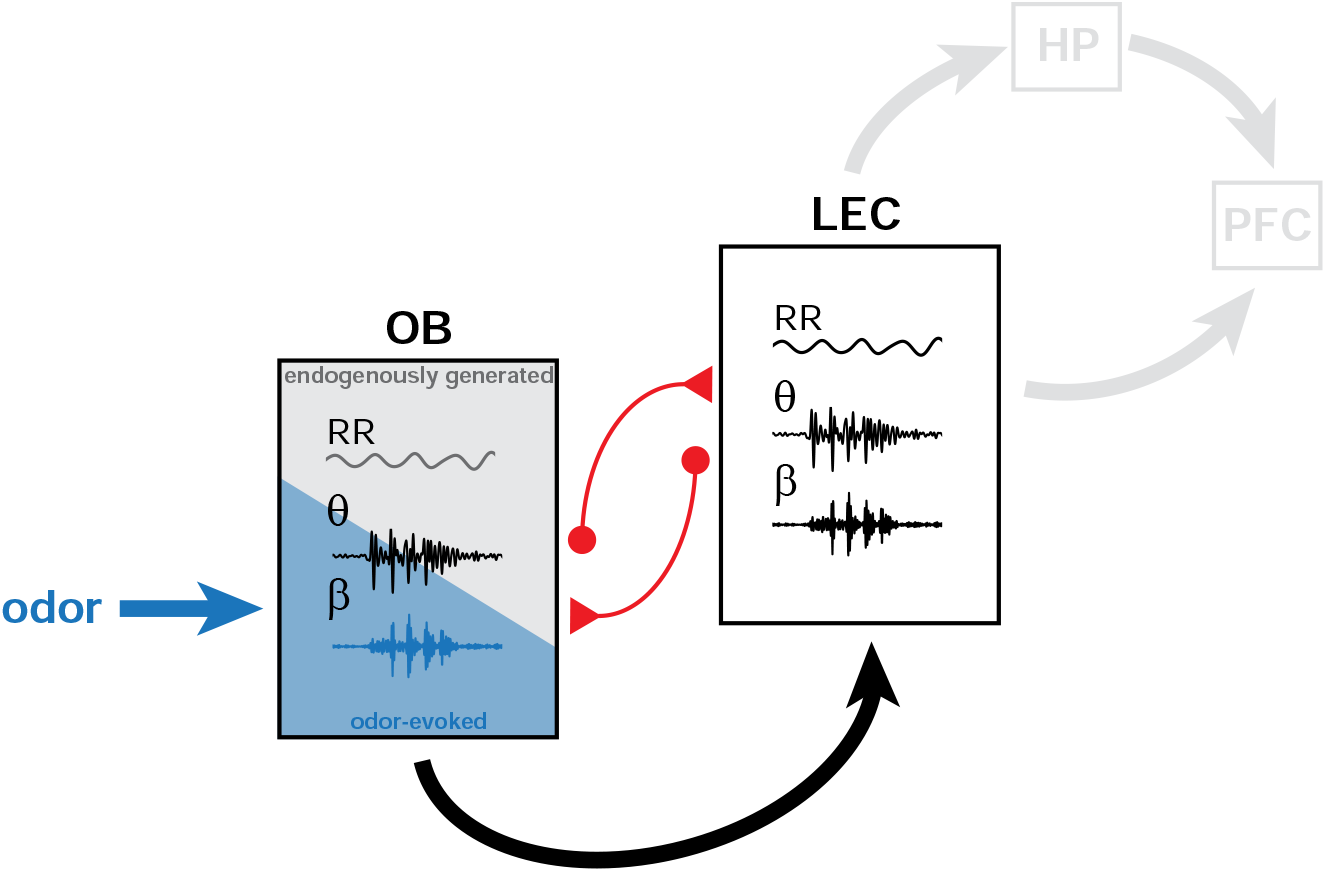
Schematic diagram of structural and functional coupling within OB-LEC networks of neonatal mice. Mutual axonal projections (red) connect neonatal OB and LEC. In OB of neonatal mice, continuous air flow-dependent RR and discontinuous MTC-driven theta bursts represent the two major patterns of oscillatory activity. They are augmented by olfactory stimuli (blue) that additionally evoke beta oscillations. OB activity boosts the oscillatory entrainment of neonatal LEC that, in turn, might drive the limbic circuits during development.

Brain development has been extensively investigated in rodents because they enable insights into a time window that remains inaccessible in humans. As altricial species, rodents are born at an immature stage of brain development. They are blind, deaf, do not whisker and have limited motor abilities during the first postnatal days. Before the onset of the ability to actively respond to sensory stimuli, coordinated activity patterns, typically characterized by rhythmic burst discharge separated by periods of quiescence, emerge endogenously. Such patterns have been described in developing somatosensory, visual and auditory systems. Their onset, properties, and underlying mechanisms are relatively well understood. For example, retinal waves emerge before the onset of light-sensitivity and vision as local patterns of coordinated activity mediated by gap junctions, cholinergic and glutamatergic circuits (53, 54). Retinal waves synaptically propagate along the visual tract to primary visual cortex (35, 55, 56) and are mandatory for the refinement of visual maps (57). Similarly, cochlear burst activity emerges before the onset of hearing as a result of coordinated firing and propagates along auditory pathways (58, 59). These cochlear bursts are crucial for the establishment of precise tonotopic maps (5, 49). The precision of whisker maps in the primary somatosensory cortex seems to be equally controlled by coordinated activity evolving during postnatal development (8, 60). In the absence of a sensory periphery with bursting activity before the onset of whisking, passive activation of whiskers is replayed within thalamo-cortical circuits and contributes to refinement of topographic maps (6).

At the same postnatal age, the olfactory system is considered to be fully mature, the sense of smell being of particular relevance for pup survival. This early maturity poses the question, whether the mechanisms of organization differ between developing olfactory pathways and other sensory systems. The present data indicate that, similar to retina or cochlea, OB generates discontinuous patterns of oscillatory activity peaking in theta frequency range. MTCs are critical mediators of theta bursts. These bursts are complemented by the continuous RR that is timed by respiration/air flow and largely independent of neuronal firing in OB. While endogenously generated discontinuous theta bursts are a common activity pattern in peripheral sensory structures, independent of system maturity (OB *versus* retina, cochlea), the continuous RR was not reported for other sensory systems. The frequency structure of network activity in neonatal OB was largely independent of the brain state. Anesthesia augmented theta but not RR power, suggesting that MTC activity is enhanced. Similarly, in adult OB, MTCs augmented odor-evoked activity, thus broadening their odor tuning under anesthesia (45, 61).

Overall, bursting OB activity during development profoundly differs from the oscillatory activity in adults. Supporting previous observations *in vitro* (62), we showed that neonatal MTCs are not only preferentially attuned to theta band activity but also contribute to its generation. In adult mice, network activity in theta band emerges from respiration-coupled sensory input in the glomerular layer (63) and MTCs are mainly involved in the generation of fast oscillatory activity in gamma band (64–67). Similarly, fast rhythms are absent in the developing OB (68) and only in the presence of odors beta band activity was induced. The protracted emergence of fast oscillations has been hypothesized to result from late integration of OB interneurons into local circuits and from age-dependent intrinsic biophysical properties of MTCs. As a consequence, it has been postulated that the developing OB encodes only first-order (e.g. odor identity) but not second-order sensory information (e.g. odor context) (45, 69).

The presence of both stimulus-related and endogenous network activity raises the question, whether and, if so, how both activity types either concurrently or independently shape the maturation of the olfactory system. Already the role of spontaneous activity endogenously generated in the sensory periphery has been subject of debate. As shown for network activity in the immature retina and cochlea, discontinuous OB bursts in neonatal mice are likely to have a permissive role in the establishment of precise connectivity that is inherent in an olfactory map (70). However, it remains unclear how the spontaneous and stimulus-evoked activities create a coherent sensory representation lacking mutual perturbations. This is a unique feature of the developing OB. Spontaneous retinal waves and cochlear bursts diminish and disappear with the onset of light sensitivity and hearing. Therefore, they do not interfere with stimulus-evoked activity. Understanding the mechanisms of theta bursts and RR during early development, as initiated in the present study, will enable us to disentangle their function(s) along the olfactory pathway.

In sharp contrast to most sensory pathways, the olfactory system bypasses a thalamic relay and directly conveys information from OB to cortical areas. Much research focused on the PIR, where the bulbar topography is largely discarded and dense inputs from OB are integrated to form odor percepts (16, 71–73). MTC axons also target LEC and in fact these axons represent the main input that rodent LEC receives (74). LEC neurons respond to odors (75, 76) and have been proposed to act as a modulator of olfactory coding through interactions with the PIR (24, 77, 78). The present results show that already at neonatal age a tight coupling links the OB with LEC. MTC axons target layer I of neonatal LEC as previously shown for adults (17, 79). These projections mediate the coupling by synchrony between the two areas as well as the early drive from OB to LEC. Remarkably, the neonatal LEC and OB show similar patterns of oscillatory activity, RR and theta bursts, albeit with lower power in LEC. The coupling by synchrony between the two areas peaked within the same frequency bands, 2-4 Hz and 4-10 Hz. Given the measures used for the assessment of synchrony it is unlikely that similarities result from volume conduction. Reflecting the more pronounced OB-to-LEC innervation as compared with *vice versa* projections, the entorhinal firing was stronger timed by the phase of RR and theta bursts in the OB than the OB firing was driven by the entorhinal activity. Interestingly, lidocaine blockade of MTC firing did not abolish the coupling between OB and LEC, suggesting that alternative pathways might contribute to the synchrony between the two areas.

While feedback projections from LEC to OB (and piriform cortex, not shown) emerge early in life, their function seems to mature postnatally to reach the anticipatory top-down modulation and optimal input discrimination that have been identified at adult stage (80). Recent findings revealed that the cellular substrate of feedforward and feedback interactions between OB, LEC and PIR of adult mice are highly complex (24). We hypothesize that, under the influence of excitatory inputs from OB, the local entorhinal circuitry is activated. The MTC target neurons in LEC are mainly glutamatergic, suggesting that coordinated OB activity causes an overall excitation in LEC that might facilitate the formation and refinement of local circuits.

Olfactory information reaches the adult HP (CA1 and dentate gyrus) via reelin-positive neurons in LEC (24, 81). Along these axonal projections, the oscillatory activity is synchronized and enables directed functional interactions between OB, LEC and HP. In turn, HP unidirectionally projects to PFC. At the functional level, the communication across areas involves oscillatory activity that temporally coordinates the neuronal assemblies. For example, respiration-related slow activity, even if subject to debate regarding its relationship to previously described slow rhythms (82), has recently been found to occur simultaneously with theta oscillations (83) and moreover entrain faster beta and gamma oscillations in LEC, HP, and PFC (46, 84, 85). Taking into account the role of HP and PFC for cognitive processing (86), the OB activity that directly entrains the limbic circuit via LEC activation might represent a powerful control mechanisms of memory and executive performance of adult (87, 88).

It is tempting to speculate about the potential functions of OB-driven entrainment of LEC during neonatal development, before the emergence of cognitive abilities. Our previous results demonstrated that LEC acts as gatekeeper of prefrontal-hippocampal interactions shortly after birth (11). Discontinuous theta bursts in LEC drive the oscillatory entrainment and time the firing of both prelimbic subdivision of PFC and CA1 area of the intermediate/ventral HP. Here we show that MTC-dependent theta activity of neonatal OB boosts RR and theta bursts in LEC. On the other hand, olfactory stimuli elicit even faster entrainment of OB-LEC circuitry, with beta band oscillations being only detectable in the presence of odors, such as octanal. An important issue that remains to be elucidated is whether specific scents that the pups naturally encounter during development, such as maternal odors, shape the network function even stronger than “artificial” odors. The effects of maternal odor on physical, neuroendocrine, and behavioral development of pups has been extensively investigated (27, 89, 90), yet very little is known about the underlying cellular and circuit mechanisms. We propose that endogenously generated and odor-evoked OB activity, especially as a result of maternal odor, might increase the level of excitability within entorhinal-prelimbic-hippocampal networks and strengthen their wiring. By these means, the olfactory system could facilitate the postnatal maturation of limbic circuitry and, ultimately, the emergence of cognitive abilities.

## MATERIALS AND METHODS

### Ethics statement

All experiments were performed in compliance with the German laws and the guidelines of the European Union for the use of animals in research and were approved by the local ethical committee (15/17).

### Experimental model and subject details

*Mice.* Timed-pregnant C57Bl/6J and Tbet-cre mice from the animal facility of the University Medical Center Hamburg-Eppendorf as well as *B6.Cg-Gt(ROSA)26Sor^tm40-1(CAG-aop3/EGFP)Hze^*/J mice (Ai40(RCL-ArchT-EGFP)-D, Jackson Laboratory, stock no: 02118), and Tbet-cre;ArchT-EGFP mice (bred by the animal facility of the University Medical Center Hamburg-Eppendorf) were housed individually in breeding cages at a 12 h light / 12 h dark cycle and fed *ad libitum.* Mouse lines used for CLARITY experiments (Tbet-cre mice, B6.Cg-*Gt(ROSA)26So^tm9(CAG-tdTomato)Hze^*/J (Ai9(RCL-tdT), Jackson Laboratory, stock no: 007909 and Tbet-cre;tdT mice) were bred in the animal facility at RWTH Aachen University under similar conditions. The day of vaginal plug detection was defined E0.5, while the day of birth was assigned as P0. Male mice underwent sensory manipulation, light stimulation, pharmacological treatment and multi-site electrophysiological recordings at P8-10. For CLARITY experiments, male and female mice were used. Genotypes were determined using genomic DNA and following primer sequence (Metabion, Planegg/Steinkirchen, Germany: for Cre in Ai40(RCL-ArchT-EGFP)-D mice: PCR forward primer 5’-ATCCGAAAAGAAAACGTTGA-3’ and reverse primer 5’-ATCCAGGTTACGGATATAGT-3’; for ROSA26-wt PCR forward primer 5’-AAAGTCGCTCTGAGTTGTTAT-3’ and reverse primer 5’-GGAGCGGGAGAAATGGATATG-3’; for GFP-tg PCR forward primer 5’-CTGGTCGAGCTGGACGGCGACG-3’ and reverse primer 5’-GTAGGTCAGGGTGGTCACGAG-3’; for Cre in Ai9(RCL-tdT) mice: forward primer 5’ CATGTCCATCAGGTTCTTGC 3’ and reverse primer 5’ AGAGAAAGCCCAGGAGCAG 3’; for tdTomato forward primer 5’ GGCATTAAAGCAGCGTATCC 3’ and reverse primer 5’ CTGTTCCTGTACGGCATGG 3’. The PCR reactions were as follows: 10 min at 95°C, 30 cycles of 45 s at 95°C, 90 s at 54°C, 90 s at 72°C, followed by a final extension step of 10 min at 72°C (Cre-tg and ROSA26-wt), 10 min at 95°C, 30 cycles of 45 s at 95°C, 90 s at 68°C, 90 s at 72°C, followed by a final extension step of 10 min at 72°C (GFP-tg). In addition to genotyping, EGFP expression in OB prior to surgery was detected using a dual fluorescent protein flashlight (Electron microscopy sciences, PA, US).

### Surgical procedures

*Surgical preparation for electrophysiology and light delivery in vitro.* For patch-clamp recordings, pups were decapitated and brains were sliced in 300 μm-thick coronal sections. Slices were incubated in oxygenated ACSF containing (in mM) 119 NaCl, 2.5 KCl, 1 NaH_2_PO_4_, 26.2 NaHCO_3_, 11 glucose, 1.3 MgSO_4_ (320 mOsm) at 37 °C. Prior to recordings, slices were maintained at room temperature and superfused with oxygenated ACSF.

*Surgical preparation for electrophysiology and light delivery in vivo.* For recordings in nonanesthetized state, 0.5% bupivacain / 1% lidocaine was locally applied on the neck muscles. For recordings under anesthesia, mice were injected i.p. with urethane (1 mg/g body weight; Sigma-Aldrich, MO, USA) prior to surgery. For both groups, under isoflurane anesthesia (induction: 5%, maintenance: 2.5%) the head of the pup was fixed into a stereotaxic apparatus as previously reported (9).

The surgery protocols are described in detail in Supplemental Information.

### Electrophysiology

*Electrophysiological recordings in vivo.* One-shank electrodes (NeuroNexus, MI, USA) with 16 recording sites were inserted into dorsal (depth 0.5-1.2 mm, angle 0°) or ventral OB (1.4-1.8 mm, angle 0°) as well as in LEC (depth: 2 mm, angle: 10° from the vertical plane). Two-shank optoelectrodes (Buzsaki16-OA16LP, NeuroNexus, MI, USA) with 8 recordings sites on each shank aligned with an optical fiber ending 40 μm above the top recording site were inserted into ventral OB. Extracellular signals were band-pass filtered (0.1 Hz – 9 kHz) and digitized (32 kHz) by a multichannel amplifier (Digital Lynx SX; Neuralynx, Bozeman, MO; USA) and Cheetah acquisition software (Neuralynx).

*Electrophysiological recordings in vitro.* Whole-cell patch-clamp recordings were performed from MTCs identified by their location in the mitral cell layer and visualized by membrane-bound EGFP. All recordings were performed at room temperature. Recording electrodes (4-9 MΩ) were filled with K-gluconate based solution containing (in mM): 130 K-gluconate, 10 HEPES, 0.5 EGTA, 4 Mg-ATP, 0.3 Na-GTP, 8 NaCl (285 mOsm, pH 7.4) and 0.5% biocytin for post-hoc morphological identification of recorded cells. Recordings were controlled with the Ephus software (91) in the MATLAB environment (The MathWorks, Inc., MA, USA).

### Morphological investigation

*CLARITY.* Brains from neonatal mice of both sexes were sliced in 1 mm- (for LEC) and 500 μm-thick (for OB) coronal sections. To maintain the structural integrity, the tissue was fixed overnight at 4°C in hydrogel fixation solution containing 4% acrylamide, 0.05% bis-acrylamide, 0.25% VA-044 Initiator, 4% PFA in PBS-/-. After polymerization and embedding the nuclear marker DRAQ5 (1:1000) was added to the samples. After washing steps, the samples were incubated for 24 h in RIMS80 containing 80 g Nycodenz, 20 mM PS, 0.1% Tween 20, and 0.01% sodium acid.

*Retrograde tracing.* For retrograde tracing, anesthetized P3-4 mice received unilateral Fluorogold (FG) (Fluorochrome, LLC, USA) injections into OB (0.8 mm anterior from the fronto-nasal suture, 0.8 mm from midline) or LEC (1 mm posterior to bregma, 5 mm from midline). After 4-5 days, pups were deeply anesthetized and perfused at P8.

All staining protocols are described in detail in Supplemental Information.

### Manipulations

*Light stimulation in vitro.* Whole-cell current-clamp recordings were performed from ArchT-EGFP expressing mitral cells in coronal slices of the neonatal Tbet-cre;ArchT mice. Yellow light pulses (595 nm) of different light intensities (0.2 – 2.6 mW) were applied to test the effect on the membrane potential.

*Light stimulation in vivo.* Trapezoid light stimulation was applied using a diode pumped solid state (DPSS) laser (Cobolt Mambo, 594 nm, Omicron, Austria), controlled by an Arduino Uno (Arduino, Italy).

*Naris occlusion.* One naris was closed using silicon adhesive (Kwik-Sil, World Precision Instruments). After a recovery period of five minutes, the recording was pursued while one naris was sealed.

*Pharmacological inactivation.* To block the firing of OB neurons, lidocaine hydrochloride 4% in 0.9% NaCl, pH 7.0 with NaOH) was slowly infused into the OB

*Odor stimulation.* An eight channel dilution olfactometer (Aurora Scientific) was used for stimulus delivery.

All manipulation protocols are described in detail in Supplemental Information.

### Quantification and statistical analysis

*Immunohistochemistry quantification.* Images were analyzed using ImageJ.

*Detection of respiration frequency.* Respiration was monitored using a piezo-electric sensor placed under the pup’s chest.

*LFP analysis.* Data were analyzed offline using custom-written scripts in the MATLAB environment (Version 9, MathWorks, Natick, MA).

For details, see Supplemental Information.

*Statistics.* Statistical analysis was performed using SPSS Statistics 22 (IBM, NY) or MATLAB. Gaussian distribution of the data was assessed using the Kolmogorov-Smirnov test. None of the data sets were normally distributed. Therefore, data were tested for significance using Wilcoxon signed-rank test (2 related samples), Wilcoxon rank-sum test (2 unrelated samples), Friedman test (>2 related samples; Wilcoxon signed-rank post hoc test with Bonferroni correction) and Kruskal-Wallis H test (>2 unrelated samples; Wilcoxon rank-sum test post hoc test with Bonferroni correction). Differences in proportions were tested using χ^2^ test. For classification of single unit responses to light stimulation, significant firing rate changes were assessed statistically using Wilcoxon signed-rank test. Data are represented as median and inter-quartile range.

## Acknowledgments

We thank A. Marquardt, C.H. Engelhardt, P. Putthoff, A. Dahlmann for excellent technical assistance, I. Hermans-Borgmeyer for help with mouse breeding, and F. Kutchera and T. Renz for optimization of the setup and olfactometer.

I. L.H.-O. acknowledges support by the ERC (ERC Consolidator Grant 681577) and by the German Research Foundation (Ha4466/10-1, SFB 936 (B5) and SPP 1665 (Ha4466/12-1)). M.S acknowledges support by the German Research Foundation (SP724/10-1). A.M.T. was enrolled in the NSF-funded exchange program of International Program for the Advancement of Neurotechnology (IPAN) of Univ. of Michigan. M.S. is a Lichtenberg-Professor of the Volkswagen Foundation.

I. L.H.-O. and M.S. are members of the FENS Kavli Network of Excellence

## Author contributions

I. L.H.-O. designed the experiments, S.G, J.K.K., H.H., K.W., D.F., A.M.T carried out the experiments, S.G., J.K.K., H.H., K.W. analyzed the data, I.L.H.-O. S.G., J.K.K. and M.S. interpreted the data and wrote the paper. All authors discussed and commented on the manuscript.

